# The formal demography of kinship III: Kinship dynamics with time-varying demographic rates

**DOI:** 10.1101/2021.03.15.435377

**Authors:** Hal Caswell, Xi Song

## Abstract

**Background:** Kinship models generally assume time-invariant demographic rates, and compute the kinship structures implied by those rates. It is important to compute the consequences of time variation in demographic rates for kinship stuctures.

**Objectives:** Our goal is to develop a matrix model for the dynamics of kinship networks subject to arbitrary temporal variation in survival, fertility, and population structure.

**Methods:** We develop a system of equations for the dynamics of the age structure of each type of kin of a Focal individual. The matrices describing survival and fertility vary with time. The initial conditions in the time-invariant model are replaced with a set of boundary conditions for initial time and initial age.

**Results:** The time-varying model maintains the network structure of the time-invariant model. In addition to the results of the time-invariant model, it provides kinship structures by period, cohort, and age. It applies equally to historical sequences of past rates and to projections of future rates. As an illustration, we analyze the kinship structure of Sweden from 1891 to 2120.

**Contribution:** The time-varying kinship model makes it possible to analyze the consequences of changing demographic rates, in the past or the future. It is easily computable, requires no simulations, and is readily extended to include additional, more distant relatives in the kinship network. The method can also be used to show the growth of families, lineages, and dynasties in populations across time and place and between social groups.

## 1 Introduction

We live in a world that changes. It changes in the past as an observed fact of history and we predict its changes in the future as scenarios. Even so, much of demography analyzes change using models that exclude change. For example, calculations of life expectancy or population growth rate based on the mortality and fertility in a consequence for a cohort living out its life under the unchanging conditions of that year. The results can then be (and are often) used to characterize change by comparing them over time or across populations.

Time-invariant models are a powerful way to explore a set of conditions by asking what would happen if those conditions were to be maintained (Keyfitz, 1972; Cohen, 1979; Caswell, 2001). The early kinship models (Lotka, 1931; Coale, 1965; Goodman, Keyfitz, and Pullum, 1974, 1975) and the more recent alternative by Caswell (2019, 2020) are all time-invariant. Every individual in the kinship network experiences a fixed set of demographic rates. The kinship structure of a focal individual describes the hypothetical situation in which she has been subject to that fixed set of rates throughout her life.

In this paper, we relax the restriction to time-invariant rates, and allow any of the rates to vary over time in any arbitrary fashion. This generalization makes two major contributions. First, it makes it possible to examine the past, by incorporating historical changes in the demographic rates, such as are accumulated in a variety of demographic databases. Second, it makes it possible to study the future in novel ways. Projections of future populations are regularly prepared by national statistical offices and international agencies. Such projections provide future scenarios of demographic rates that can be incorporated directly into our model. The properties of the kinship network are now readily obtained as outputs of routine population projections.

Goodman, Keyfitz, and Pullum (1974, 1975) analyzed kinship structures using a system of integral equations to calculate the expected number of kin, of each type, associated with an individual of a given age. This classical approach is challenging to implement, gives only the numbers, not age distributions of kin, and is difficult to extend beyond the simplest case. A new approach was introduced by Caswell (2019, 2020), using matrix operations to project the population of each type of kin from one age to the next and account for the production of new kin of one type (e.g., nieces) by the reproduction of a different type (e.g., sisters). The first of these papers provided the general theory for the dynamics of the age structure of any kind of kin; the second extended the theory to multistate models that project the age × stage structure of all types of kin and applied the theory to analyze the age × parity structure of kin. The matrix approach simplifies model notation, facilitates computation, and uses matrix algebra to link kinship relations with population dynamics.

We will begin by reviewing briefly the time-invariant kinship model, and use that framework to develop the time-varying model. We will then use the model to analyze the past and future of kinship structure in Sweden by coupling a long sequence of historical data (1891– 2018) to a long projection of future demographic rates (2018–2120). The results shed light on current and future kinship structures in Sweden as a result of demographic transitions in fertility and mortality over the past two centuries, and provide an example for analysis of other countries.

### 1.1 Kinship with time-invariant rates

#### Notation

In what follows, matrices are denoted by boldface upper case letters (e.g., **U**) and vectors by boldface lower case letters (e.g., **a**). Vectors are column vectors; the vector **x**^⊤^ is the transpose of **x**. The symbol ||**x**|| denotes the 1-norm of **x**; i.e. the sum of the absolute values of the entries of **x**. The vector **e**_*i*_ is the ith unit vector; i.e., a vector of zeros with a 1 in the *i*th entry. The dimension of **e**_*i*_ will be specified if it is not clear from the context. We will sometimes use Matlab notation in which **F**(*i*,:) and **F**(:,*j*) denote the *i*th row and the *j*th column of **F**, respectively. The operator ∘ denotes the Hadamard, or element-by-element product.

To review briefly here, kinship is defined relative to a focal individual, who we name Focal. The kin, of any type, of Focal are a population, whose structure is governed by births of new members and deaths and aging of current members. Each type of kin in the network of Figure 1 is identified with a letter; e.g., **a**(*x*) is the age structure vector of the daughters of Focal at Focal’s age *x*. We reserve the symbol **k** as a generic kin age structure vector.

**Figure 1:**
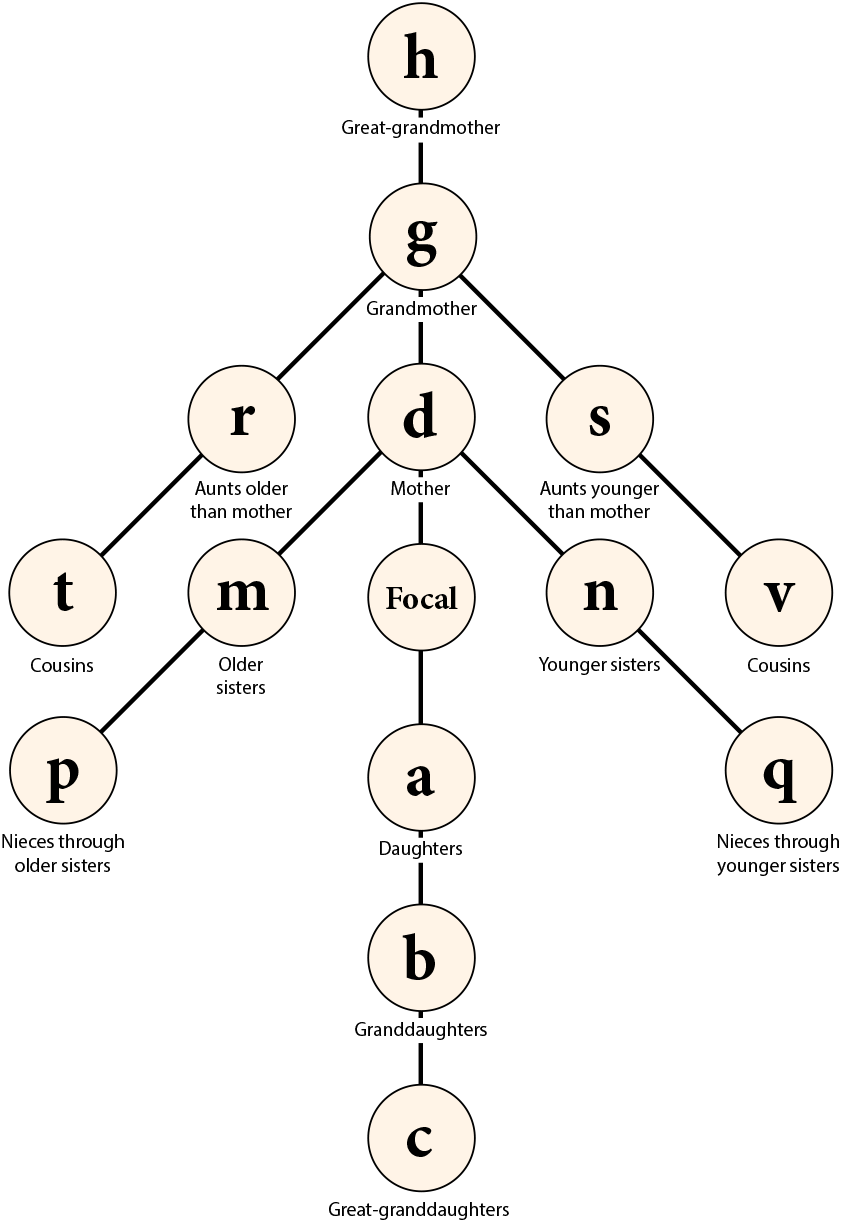
The kinship network of Focal. The population of each type of kin is denoted by a letter (**a**, **b**, etc.) which appears as an age structure vector in the kinship model. From Caswell (2019) under a CC Attribution license.

We write **k**(*x*) as the age structure vector for kin of type **k** at age *x* of Focal. The kin **k**(*x* + 1) at age *x* + 1 consists of survivors of the kin at age *x* and new kin:

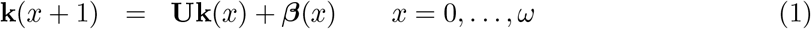

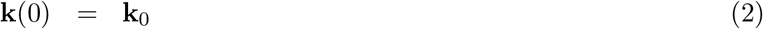

where **U** is the survival matrix and ***β***(*x*) is the age structure vector of new kin arriving between *x* and *x* + 1. For some types of kin, ***β***(*x*) = **0**, because no new kin are possible. For example, Focal cannot gain any new older sisters. For other types of kin,

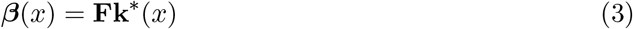

where **F** is the fertility matrix and **k**\* is some other type of kin, whose offspring are kin of type **k**. For example, new granddaughters (**b**) are produced by the reproduction of daughters (**a**). The dynamics of granddaughters are then

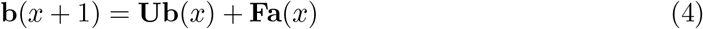

Because the model is a dynamic system, it requires an initial condition **k**(0) = **k**_0_, which gives the age structure of the kin at the birth of Focal. For some types of kin,

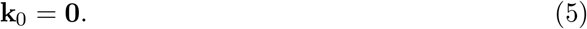

For some types of kin, the initial condition is zero (e.g., we know that Focal has no daughters at the time of her birth). For other types of kin, the initial condition is calculated as a mixture over the age distribution ***π*** of mothers at the birth of Focal (e.g., the older sisters of Focal at her birth are the daughters of Focal’s mother at her age at the birth of Focal).

The time-invariant kinship model is described in detail in Caswell (2019) with a step-by-step construction of the subsidy terms and the initial conditions for all the types of kin in Figure 1. Appendix A summarizes these calculations.

## 2 Kinship with time-varying rates

Having carefully specified the model, we are in a position to replace the time-invariant rates by allowing survival and fertility to vary over a sequence of *T* years:

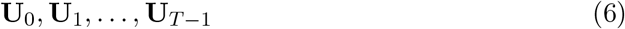

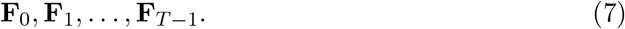

The kin population now depends on both the age of Focal and time, so we write

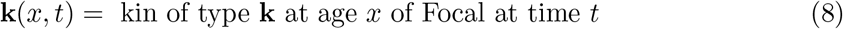

The dynamics are given by

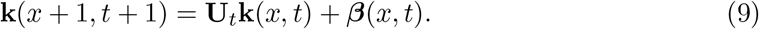

Our goal is to calculate these vectors for all the types of kin in the kinship network, over some specified time span for which we have demographic rates.

As before, the subsidy vector ***β***(*x, t*) takes one of two forms.

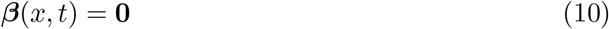

if there are no new kin of this type (e.g., older sisters of Focal). Or,

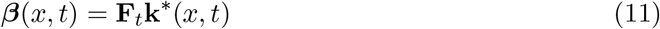

which applies the fertility at time *t* to the age structure vector of the kin that provides the subsidy. For example, the dynamics of granddaughters, given in the time-invariant case by (4), now become

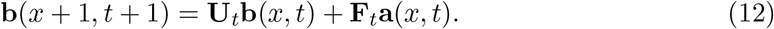

The distribution ***π*** of the ages of mothers at the birth of their children plays an important role in the calculations. In the time-invariant case, this was calculated from the stable population **w** implied by the time-invariant rates **U** and **F**. Because the rates are changing over time, the stable population associated with those rates is a poor choice of a basis for this calculation. Now we require a time-dependent distribution ***π***(*t*). Let **z**(*t*) be the age structure vector of the population (n.b., the entire population, not just one type of kin) at time *t*, given by

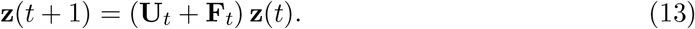

Then the vector

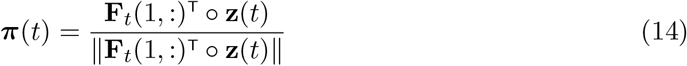

gives the distribution of ages of mothers reproducing according to **F**_*t*_ in the population at time *t*.

### 2.1 Boundary conditions

The two-dimensional dependence on age and time means that the initial condition **k**(0) in the time-invariant model is replaced by a set of boundary conditions. The dynamics in (9) requires us to specify the complete age vector at time *t* = 0,

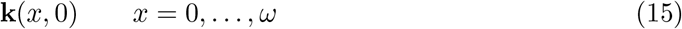

and the initial age vector at each time,

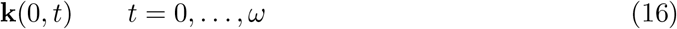

In the domain shown in Figure 2, with time on the abscissa and age on the ordinate, these boundaries correspond to the bottom and the left margins.

**Figure 2:**
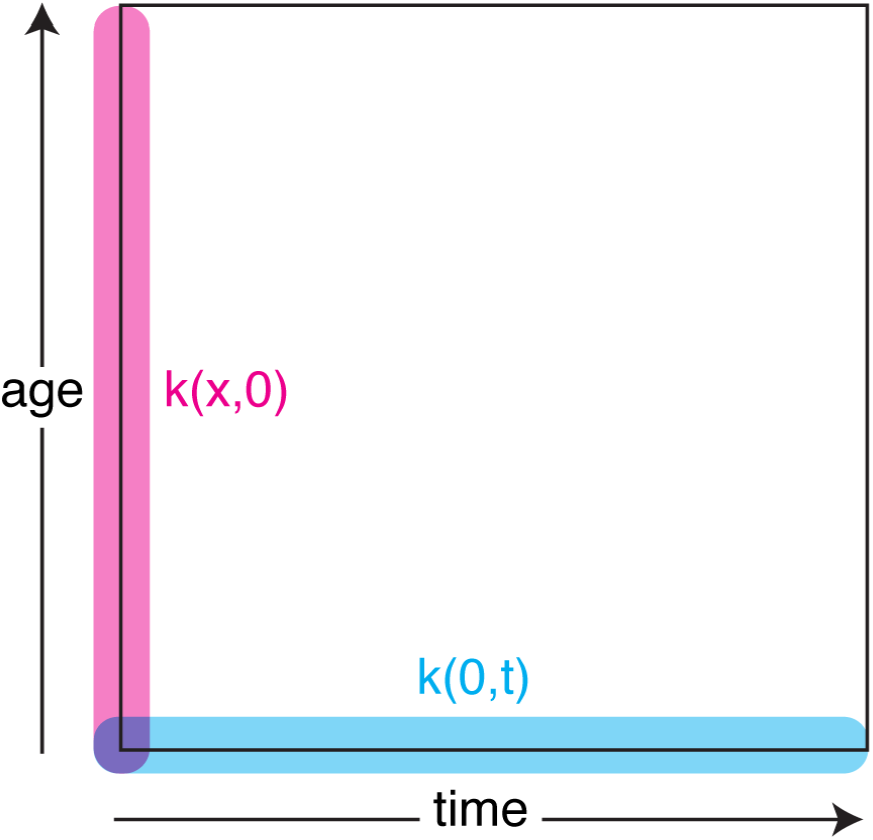
Boundary conditions. The figure contains ages from 0 to *ω* and times from 0 to *T*. The boundary conditions correspond to *k*(*x*, 0) for all *x* from 0 to *ω* and **k**(0, *t*) for all *t* from 0 to *T*.

As always, the boundary conditions require extra information (or assumptions) from outside the system. We use the following procedures

1. To calculate the time boundary **k**(*x*, 0), we suppose that the earliest available rates **U**_0_ and **F**_0_ have been operating for a long time. We use the time-invariant calculations based on these rates to generate, exactly as in Table 1 of Caswell (2019), the vectors **k**(*x*, 0) for all *x*.
2. To calculate the age boundary **k**(0, *t*), we again encounter two possibilities. If Focal has no kin of this type at her birth, then

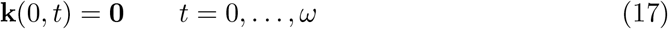 If Focal has possible kin of this type at birth, then the age boundary is calculated as

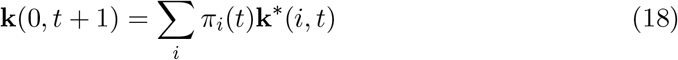

where **k**\* is an appropriate other kind of kin and *π_i_*(*t*) is the proportion of mothers who reproduce at age *i* at time *t*. These age boundaries will be presented for each type of kin in Section 2.3 and summarized in Table 1.

**Table 1:** Summary of the components of the kin model given in equation (9). For each type of kin, the relevant age boundary condition, survival dynamics, and reproductive subsidy are shown. Compare with the time-invariant model in Appendix A.

| Kin | age boundary | survival | subsidy $\beta(x)$ |
| --- | --- | --- | --- |
| $\mathbf{a}(x, t)$ daughters | $\mathbf{0}$ | $\mathbf{U}_t \mathbf{a}(x, t)$ | $\mathbf{F}_t \mathbf{e}_x$ |
| $\mathbf{b}(x, t)$ granddaughters | $\mathbf{0}$ | $\mathbf{U}_t \mathbf{b}(x, t)$ | $\mathbf{F}_t \mathbf{a}(x, t)$ |
| $\mathbf{c}(x, t)$ great-granddaughters | $\mathbf{0}$ | $\mathbf{U}_t \mathbf{c}(x, t)$ | $\mathbf{F}_t \mathbf{b}(x, t)$ |
| $\mathbf{d}(x, t)$ mothers | $\pi(t)$ | $\mathbf{U}_t \mathbf{d}(x, t)$ | $\mathbf{0}$ |
| $\mathbf{g}(x, t)$ grandmothers | $\sum_i \pi_i(t) \mathbf{d}(i, t)$ | $\mathbf{U}_t \mathbf{g}(x, t)$ | $\mathbf{0}$ |
| $\mathbf{h}(x, t)$ great-grandmothers | $\sum_i \pi_i(t) \mathbf{g}(i, t)$ | $\mathbf{U}_t \mathbf{h}(x, t)$ | $\mathbf{0}$ |
| $\mathbf{m}(x, t)$ older sisters | $\sum_i \pi_i(t) \mathbf{a}(i, t)$ | $\mathbf{U}_t \mathbf{m}(x, t)$ | $\mathbf{0}$ |
| $\mathbf{n}(x, t)$ younger sisters | $\mathbf{0}$ | $\mathbf{U}_t \mathbf{n}(x, t)$ | $\mathbf{F}_t \mathbf{d}(x, t)$ |
| $\mathbf{p}(x, t)$ nieces via older sisters | $\sum_i \pi_i(t) \mathbf{b}(i, t)$ | $\mathbf{U}_t \mathbf{p}(x, t)$ | $\mathbf{F}_t \mathbf{m}(x, t)$ |
| $\mathbf{q}(x, t)$ nieces via younger sisters | $\mathbf{0}$ | $\mathbf{U}_t \mathbf{q}(x, t)$ | $\mathbf{F}_t \mathbf{n}(x, t)$ |
| $\mathbf{r}(x, t)$ aunts older than mother | $\sum_i \pi_i(t) \mathbf{m}(i, t)$ | $\mathbf{U}_t \mathbf{r}(x, t)$ | $\mathbf{0}$ |
| $\mathbf{s}(x, t)$ aunts younger than mother | $\sum_i \pi_i(t) \mathbf{n}(i, t)$ | $\mathbf{U}_t \mathbf{s}(x, t)$ | $\mathbf{F}_t \mathbf{g}(x, t)$ |
| $\mathbf{t}(x, t)$ cousins; aunts older than mother | $\sum_i \pi_i(t) \mathbf{p}(i, t)$ | $\mathbf{U}_t \mathbf{t}(x, t)$ | $\mathbf{F}_t \mathbf{r}(x, t)$ |
| $\mathbf{v}(x, t)$ cousins; aunts younger than mother | $\sum_i \pi_i(t) \mathbf{q}(i, t)$ | $\mathbf{U}_t \mathbf{v}(x, t)$ | $\mathbf{F}_t \mathbf{s}(x, t)$ |

### 2.2 Period, cohort, and age calculations

Including time variation in the demographic rates invites the calculation of period-specific, cohort-specific, and age-specific results, as shown in Figure 3.

**Figure 3:**
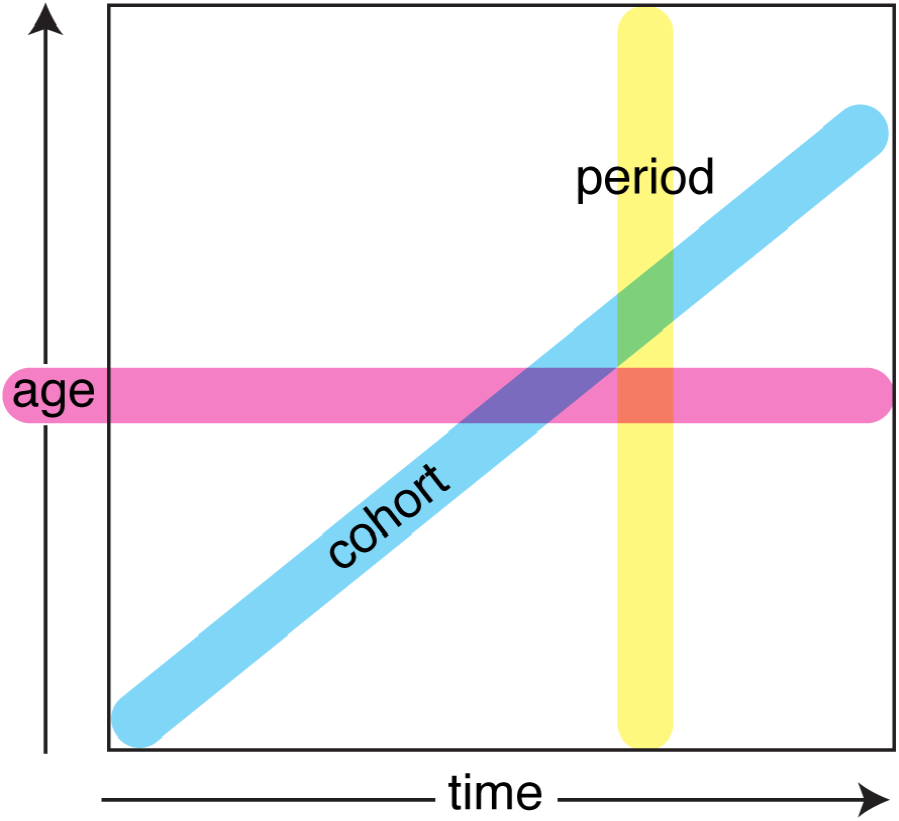
The period, cohort, and age dimensions of kinship development, within the age×time domain shown in Figure 2

#### Period results

The kinship structure vector, observed at a specified period 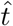, as a function of the age of Focal:

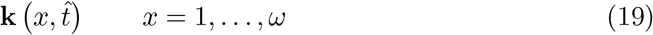

#### Cohort results

The kinship structure of Focal as a member of a cohort starting at a specified time *t*_0_ and age *x*_0_:

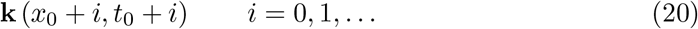

For birth cohorts, *x*_0_ = 0. The cohort can be followed until either *x*_0_ + *i* = *ω* or *t*_0_ + *i* = *T*, whichever comes first.

#### Age results

The kinship structure vector of Focal at a specified age 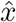, observed as a function of time:

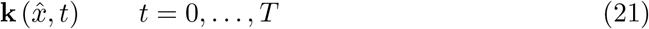

### 2.3 Dynamics of the kinship network

In this section, we derive the dynamics of each of the types of kin shown in Figure 1.^1^ The results are summarized in Table 1.

#### 2.3.1 Daughters and descendants

Each type of descendent depends on the reproduction of another type of descendent, or of Focal herself.

##### a(*x, t*) = daughters of Focal

Daughters are the result of the reproduction of Focal. Since Focal is assumed to be alive at age x, the subsidy vector is ***β***(*x, t*) = **F**_*t***e**_*x*__, where **e**_*x*_ is the unit vector for age *x*. Because we may be sure that Focal has no daughters when she is born, the initial condition is **a**(0, *t*) = **0**. Thus

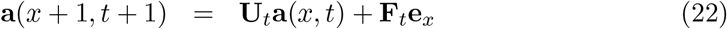

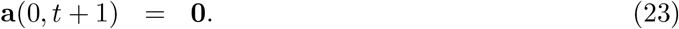

##### b(*x, t*) = granddaughters of Focal

Granddaughters are the children of the daughters of Focal. At age x of Focal, these daughters have age distribution **a**(*x, t*), so ***β***(*x, t*) = **F**_*t*_**a**(*x, t*). Because Focal has no granddaughters at birth, the initial condition is **0**;

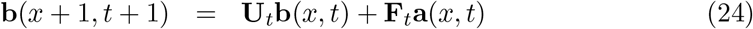

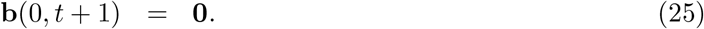

##### c(*x, t*) = great-granddaughters of Focal

Similarly, great-granddaughters are the result of reproduction by the granddaughters of Focal, with an initial condition of **0**.

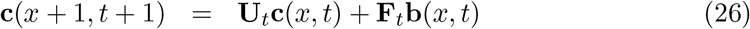

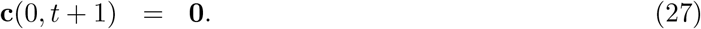

The extension to arbitrary levels of direct descendants is obvious. Let **k**_*n*_, in this case, be the age distribution of descendants of level *n*, where *n* = 1 denotes children. Then

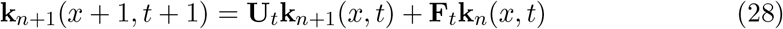

with the initial condition **k**_*n*+1_(0, *t* + 1) = **0**. The distribution of descendants in any future generation can be estimated on the basis of the recursive form of the two-generation model in (28).

#### 2.3.2 Mothers and ancestors

The surviving mothers and other direct ancestors of Focal depend on the age of those ancestors at the time of the birth of Focal.

##### d(*x, t*) = mothers of Focal

The population of mothers of focal consists of at most a single individual (step-mothers are not considered here). This population has an expected age distribution, and is subject to survival according to **U**_*t*_. No new mothers arrive after Focal’s birth, so the subsidy term is ***β***(*x, t*) = **0**.

At the time of Focal’s birth, she has exactly one mother, but we do not know her age. Hence the initial age distribution **d**(0, *t*) of mothers is a mixture of unit vectors **e**_*i*_; the mixing distribution is the distribution ***π***(*t*) of ages of mothers given by equation (14). Thus,

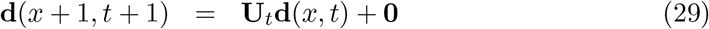

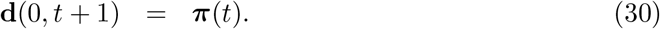

##### g(*x*) = grandmothers of Focal

The grandmothers of Focal are the mothers of the mother of Focal. No new grandmothers appear after Focal is born, so once again the subsidy term ***β***(*x, t*) = **0**. The age distribution of grandmothers at the birth of Focal is the age distribution of the mothers of Focal’s mother, at the age of Focal’s mother when Focal is born. The age of Focal’s mother at Focal’s birth is unknown, so the initial age distribution of grandmothers is a mixture of the age distributions **d**(*x, t*) of mothers, with mixing distribution ***π***:

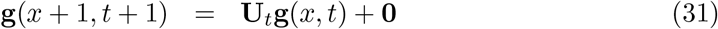

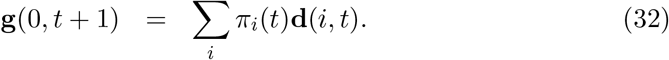

##### h(*x, t*) = great-grandmothers of Focal

Again, the subsidy term is ***β***(*x, t*) = **0**. The initial condition is a mixture of the age distributions of the grandmothers of Focal’s mother, with mixing distribution ***π***:

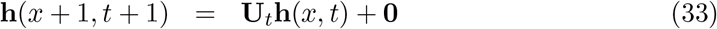

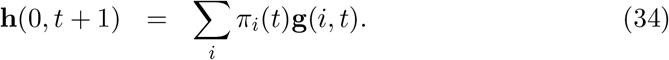

The extension to arbitrary levels of direct ancestry is clear. Let **k**_*n*_ be, in this case, the age distribution of ancestors of level *n*, where *n* = 1 denotes mothers. Then the dynamics and initial conditions are

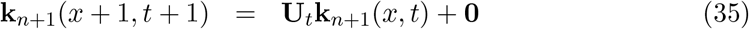

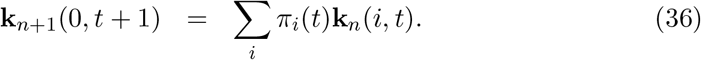

Note that, because Focal has at most one mother, grandmother, etc., the expected number of mothers, grandmothers, etc. is also the probability of having a living mother, grandmother, etc.

#### 2.3.3 Sisters and nieces

The sisters of Focal, and their children, who are the nieces of Focal, form the first set of side branches in the kinship network of Figure 1. Following Goodman, Keyfitz, and Pullum (1974), we divide the sisters of Focal into older and younger sisters because they follow different dynamics.

##### m(*x, t*) = older sisters of Focal

Once Focal is born, she can accumulate no more older sisters, so the subsidy term is ***β***(*x, t*) = **0**. At Focal’s birth, her older sisters are the children **a**(*i, t*) of the mother of Focal at the age i of Focal’s mother. This age is unknown, so the initial condition **m**(0, *t*) is a mixture of the age distributions of children with the mixing distribution ***π***(*t*).

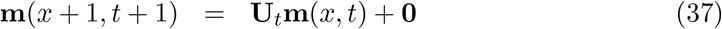

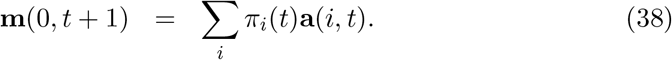

##### n(*x, t*) = younger sisters of Focal

Focal has no younger sisters when she is born, so the initial condition is **n**(0, *t*) = **0**. Younger sisters are produced by reproduction of Focal’s mother, so the subsidy term is the reproduction of the mothers at age *x* of Focal.

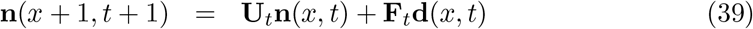

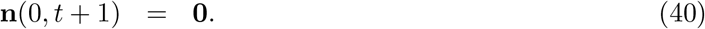

##### p(*x, t*) = nieces through older sisters of Focal

At the birth of Focal, these nieces are the granddaughters of the mother of Focal, so the initial condition is mixture of granddaughters with mixing distribution ***π***(*t*). New nieces through older sisters are the result of reproduction by the older sisters, at age *x*, of Focal.

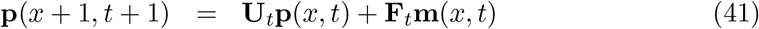

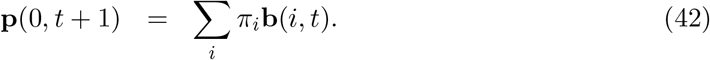

##### q(*x, t*) = nieces through younger sisters of Focal

At the birth of Focal she has no younger sisters, and hence has no nieces through these sisters. Thus the initial condition is **q**(0, *t*) = **0**. New nieces are produced by reproduction of the younger sisters of Focal.

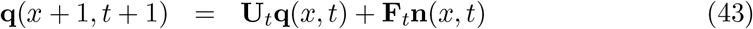

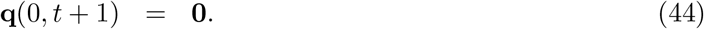

#### 2.3.4 Aunts and cousins

Aunts and cousins form another level of side branching on the kinship network; their dynamics follow the same principles as those for sisters and nieces.

##### r(*x, t*) = aunts older than mother of Focal

These aunts are the older sisters of the mother of Focal. Once Focal is born, her mother accumulates no new older sisters, so the subsidy term is ***β***(*x, t*) = **0**. The initial age distribution of these aunts, at the birth of Focal, is a mixture of the age distributions **m**(*x, t*) of older sisters, with mixing distribution ***π***(*t*)

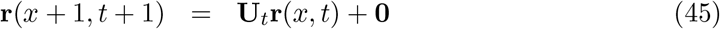

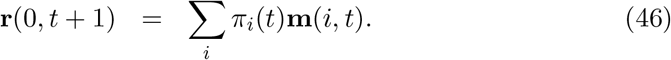

##### s(*x, t*) = aunts younger than mother of Focal

These aunts are the younger sisters of the mother of Focal. These aunts are the children of the grandmother of Focal, and thus the subsidy term comes from reproduction by the grandmothers of Focal. The initial age distribution of these aunts, at the birth of Focal, is a mixture of the age distributions **n**(*x, t*) of younger sisters, with mixing distribution ***π***(*t*).

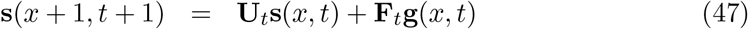

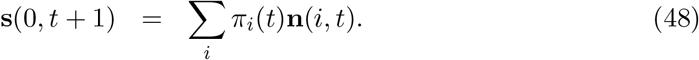

##### t(*x, t*) = cousins from aunts older than mother of Focal

These are the children of the older sisters of the mother of Focal, and thus the nieces of the mother of Focal through her older sisters. The subsidy term comes from reproduction by the older sisters of the mother of Focal.The initial condition is a mixture of the age distributions of nieces through older sisters, with mixing distribution ***π***(*t*),

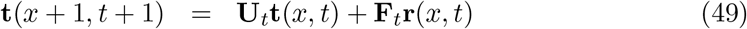

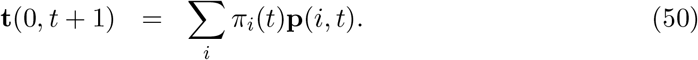

Let there be no confusion between the vector **t** giving the age structure of cousins and the scalar *t* indexing time.

##### v(*x, t*) = cousins from aunts younger than mother of Focal

These are the nieces of the mother of Focal through her younger sisters. The subsidy term comes from reproduction by the younger sisters of the mother of Focal. The initial condition is a mixture of the age distributions of nieces through younger sisters, with mixing distribution ***π***(*t*).

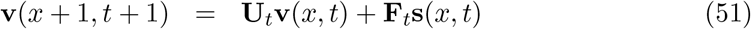

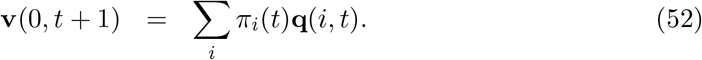

## 3 Kinship in Sweden: the past and the future

As an example combining the history of the past and projection of the future, we have analyzed the kinship dynamics of Sweden from 1891 to 2021.

We obtained mortality and fertility schedules for Swedish females from 1891 to 2018 (the past) from the Human Mortality and Human Fertility databases (Human Fertility Database, 2020; Human Mortality Database, 2020). We obtained mortality and fertility schedules from 2018 to 2120 (the future) from the population projections of the Swedish Statistical Office (Statistics Sweden; Sveriges Officiella Statistik)^2^.

The HMD and HFD data include ages 0-110. The projections by Statistics Sweden include ages 0-105. To obtain a consistent matrix dimension, we incorporated five additional age classes in the projected schedules by setting survival for ages 106-110 equal to the survival of age 105. Age-specific fertility for ages 106-110 was, of course, set to 0. Given this modification, we were able to splice together the past and the future.

The resulting trajectories of life expectancy and total fertility rate (TFR) over the entire 229 year period are shown in Figure 4. From 1891-2018, Sweden experienced a typical gradual increase in life expectancy at birth, from about 50 years to almost 85 years. Fertility declined precipitously from a TFR of just over 4 in 1891 to less than 2 by the late 1930s. A postwar baby boom increased TFR to just over 2 from 1945 to 1965, and TFR has fluctuated around 2 since then.

**Figure 4:**
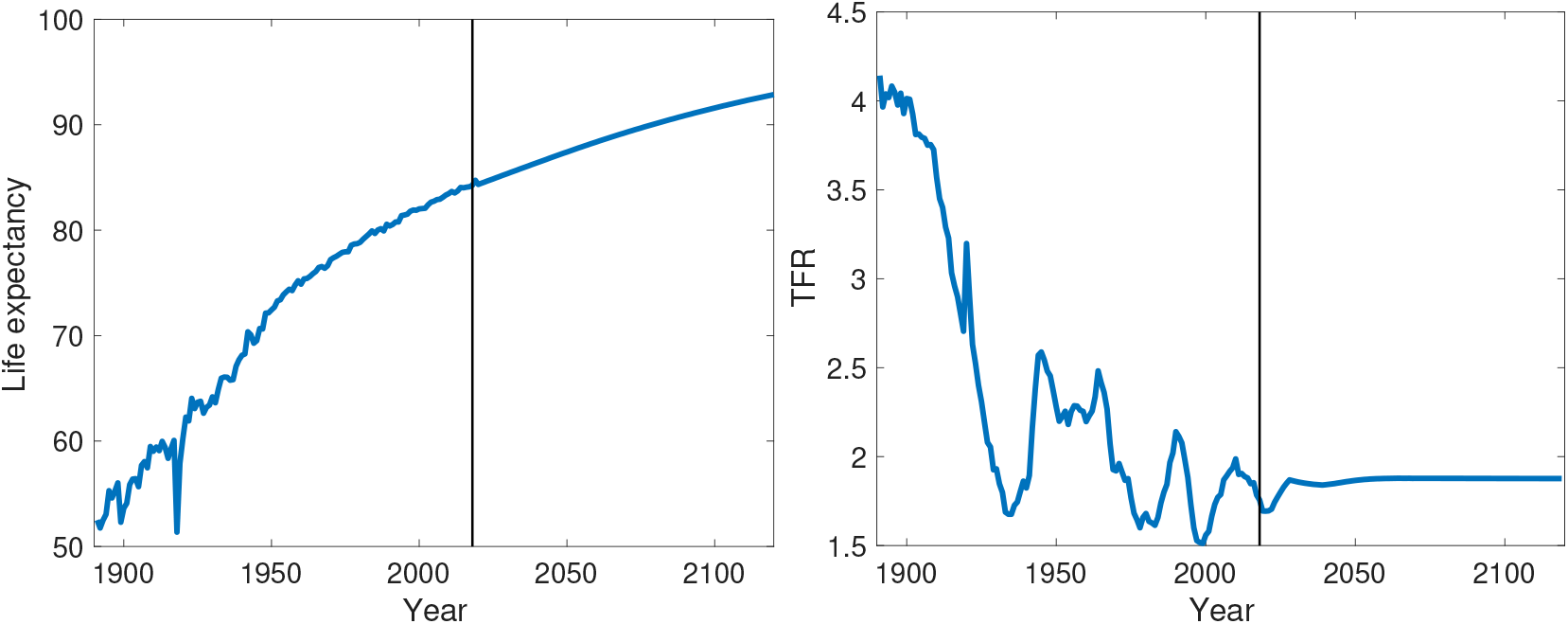
The life expectancy at birth (left) and the total fertility rate TFR (right) for Swedish females. The vertical line distinguishes past values (1891–2018) from future projections (2019–2120).

The future projected by Statistics Sweden is much calmer than the past, especially for fertility. It is interesting to note that the 1918 influenza pandemic created a noticeable dip in life expectancy, the eventual effects of the COVID-19 pandemic were not (and could not have been) projected.

As discussed in previous demographic work, we expect that the increased survival and reduced fertility over time will have opposing effects on the kinship network: reduced fertility means that fewer kin will be created, but increased survival means that fewer will be lost to mortality (Bengtson, 2001; Mare, 2011; Seltzer, 2019; Uhlenberg, 1996, 2009).

The results of the analysis of kinship in Sweden are presented in a series of figures (4-A to 4-E). These graphs combine the historical record of the past (1891-2018) and projections of the future (2019-2120). The 14 kin types in Figure 1 have been reduced to 10 by combining older and younger sisters into a single category of sisters, and similarly for nieces through older and younger sisters, aunts older and younger than mother, and cousins through older and younger aunts.

**Figure 4-A:**
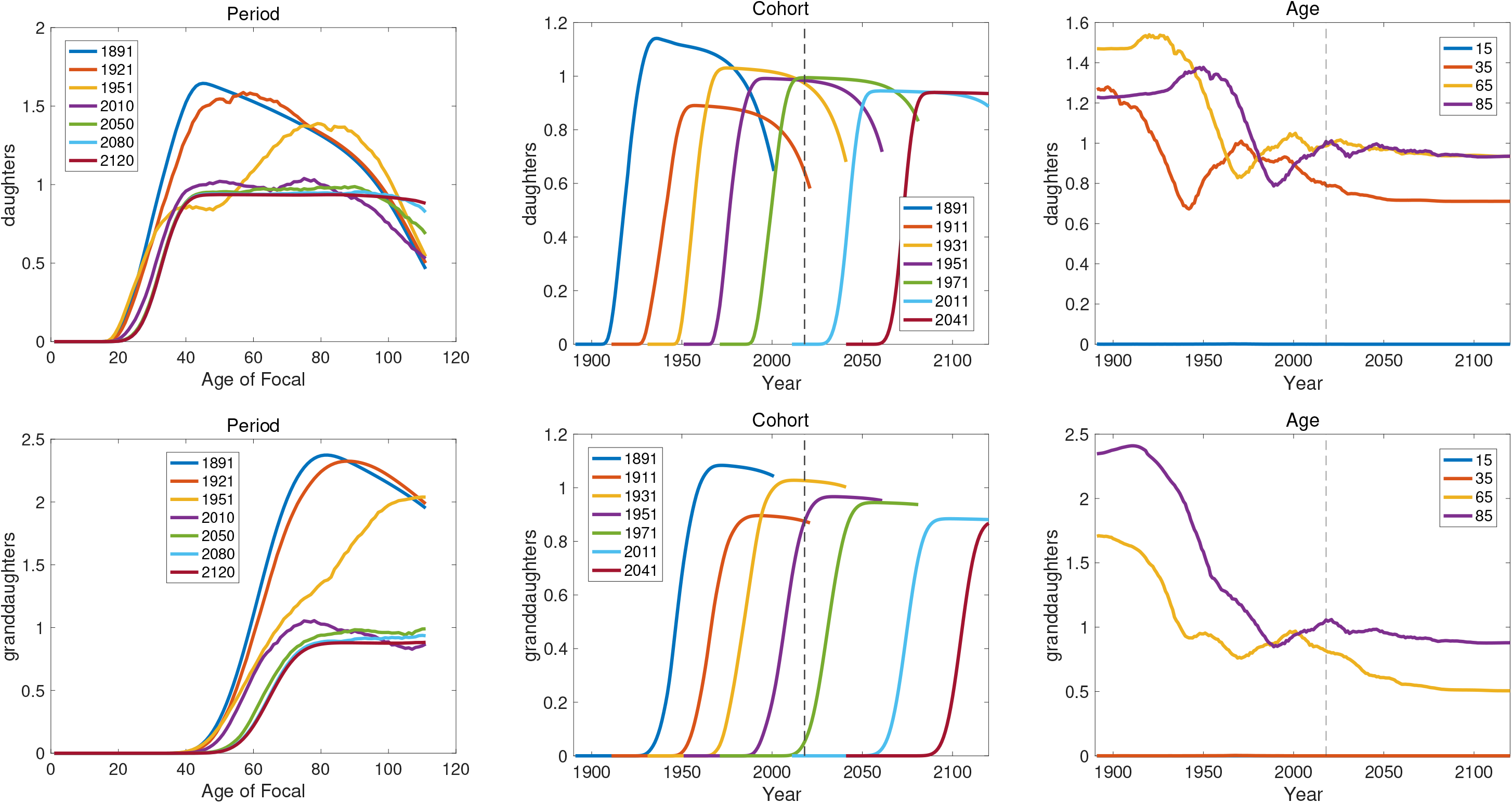
Period, cohort, and age results for the numbers of daughters and granddaughters of Focal; data from Sweden. Period kin are shown as a function of the age of Focal, for selected years from 1891 to 2120. Numbers of kin for the cohort of which Focal is a member are shown for cohorts starting in selected years from 1891 to 2041. Numbers of kin for selected ages of Focal are shown over years from 1891 to 2120. The vertical line at year 2018 marks the division between rates calculated from observations of the past and rates projected into the future.

**Figure 4-B:**
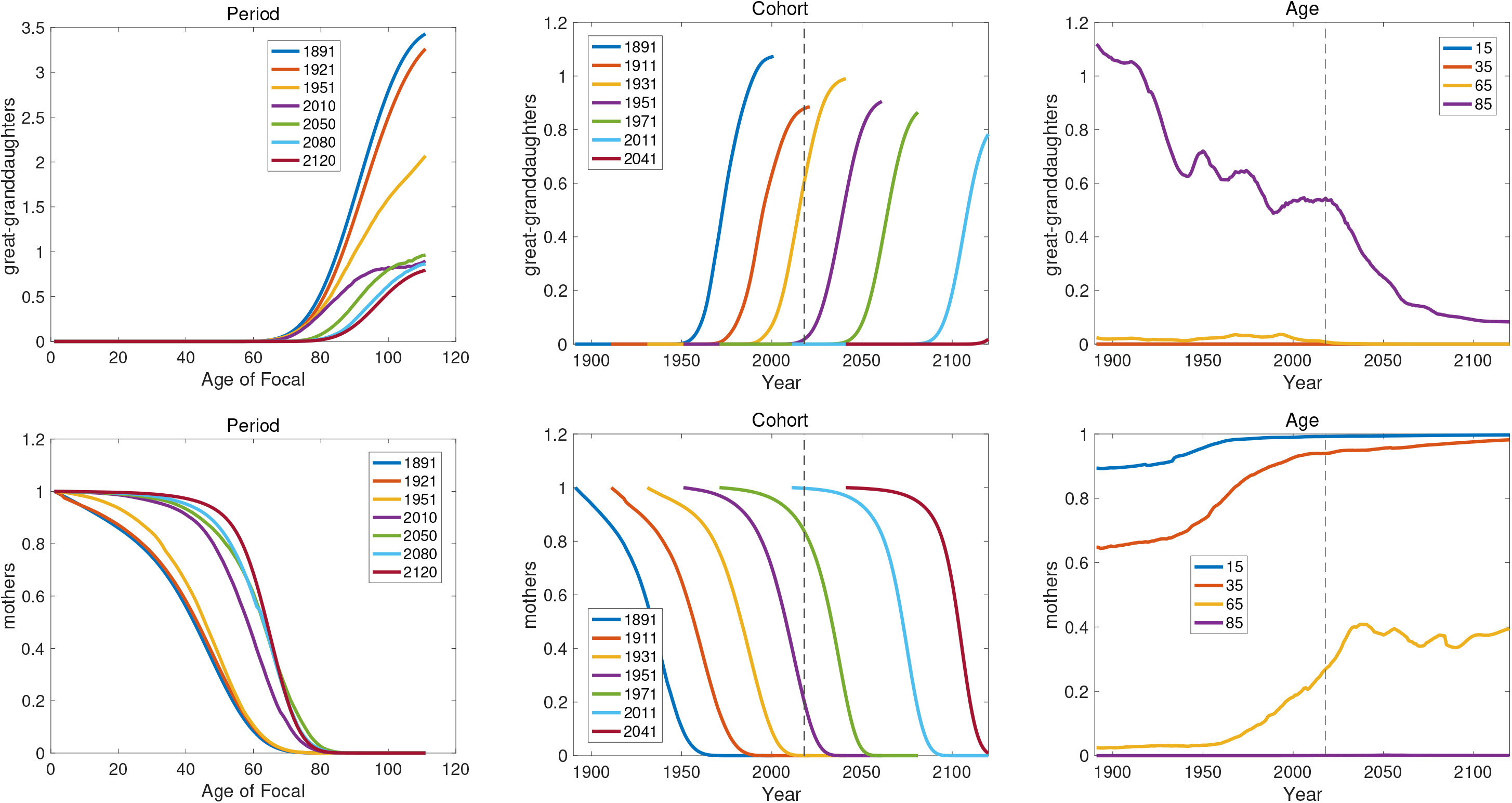
Period, cohort, and age results for the numbers of great-granddaughters and mothers of Focal; data from Sweden.

**Figure 4-C:**
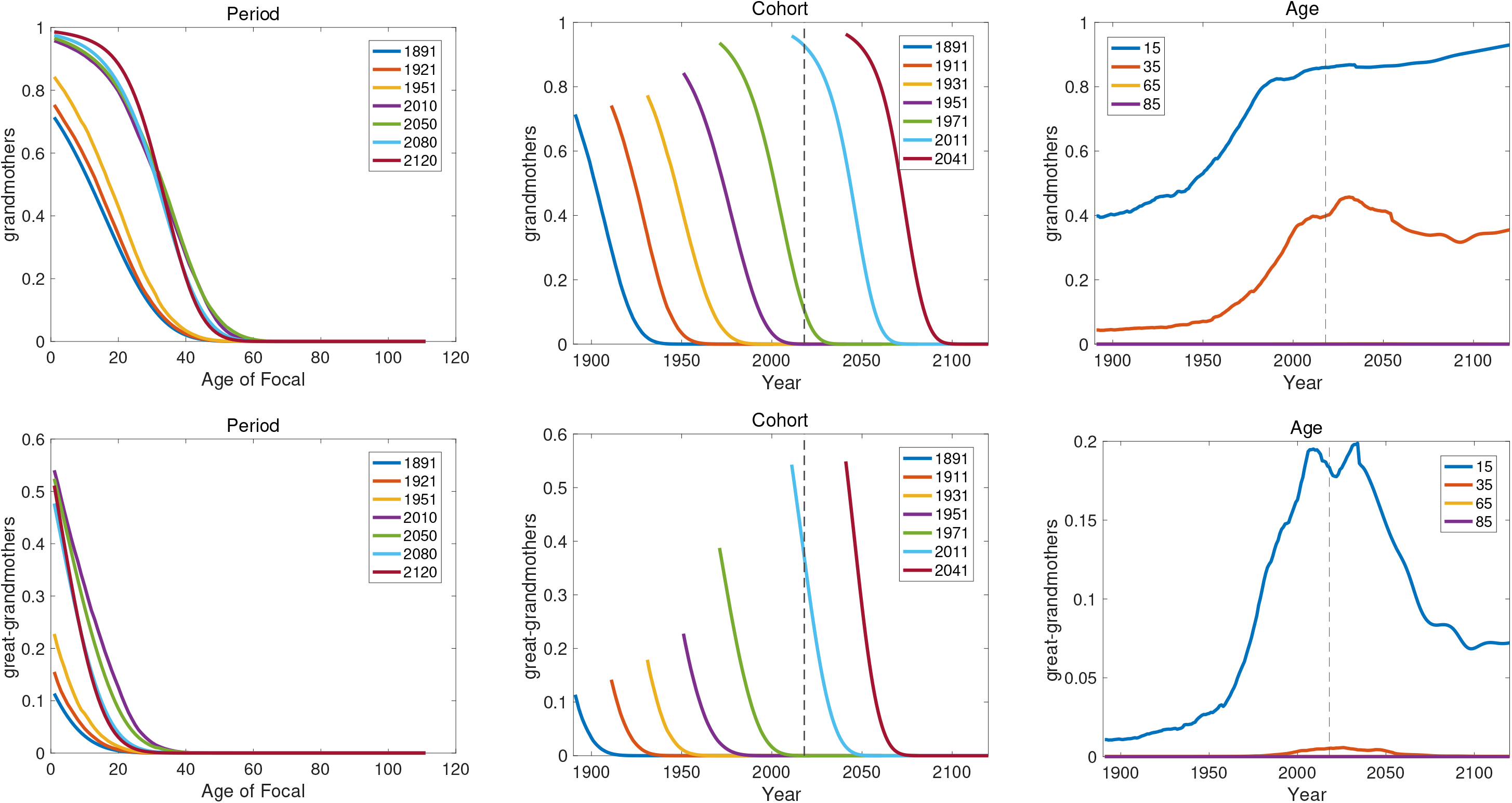
Period, cohort, and age results for the numbers of great-grandmothers and great-grandmothers of Focal; data from Sweden.

**Figure 4-D:**
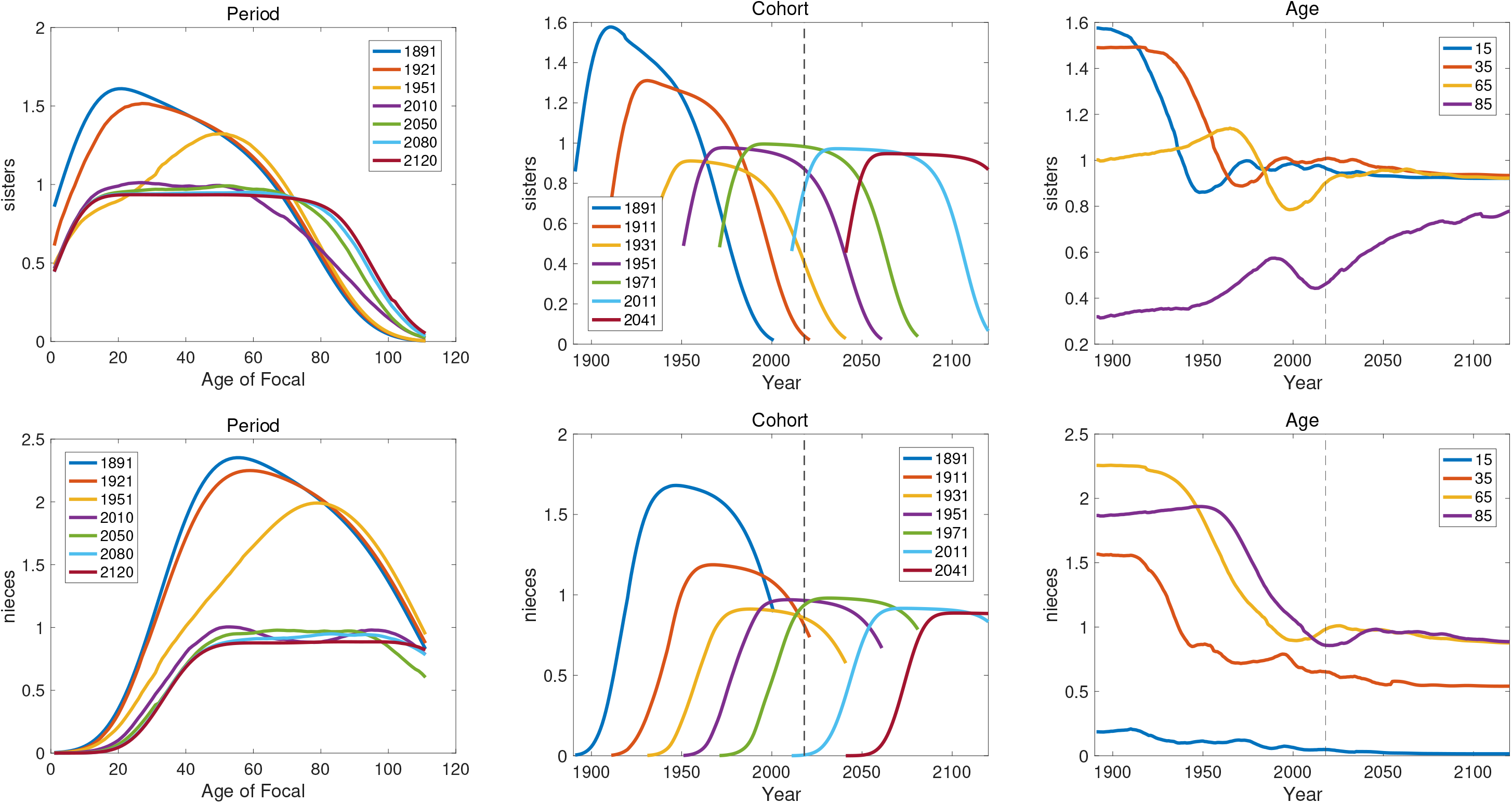
Period, cohort, and age results for the numbers of sisters and nieces of Focal; data from Sweden.Older and younger sisters, and nieces through older and younger sisters as shown in Figure 1, have been combined.

**Figure 4-E:**
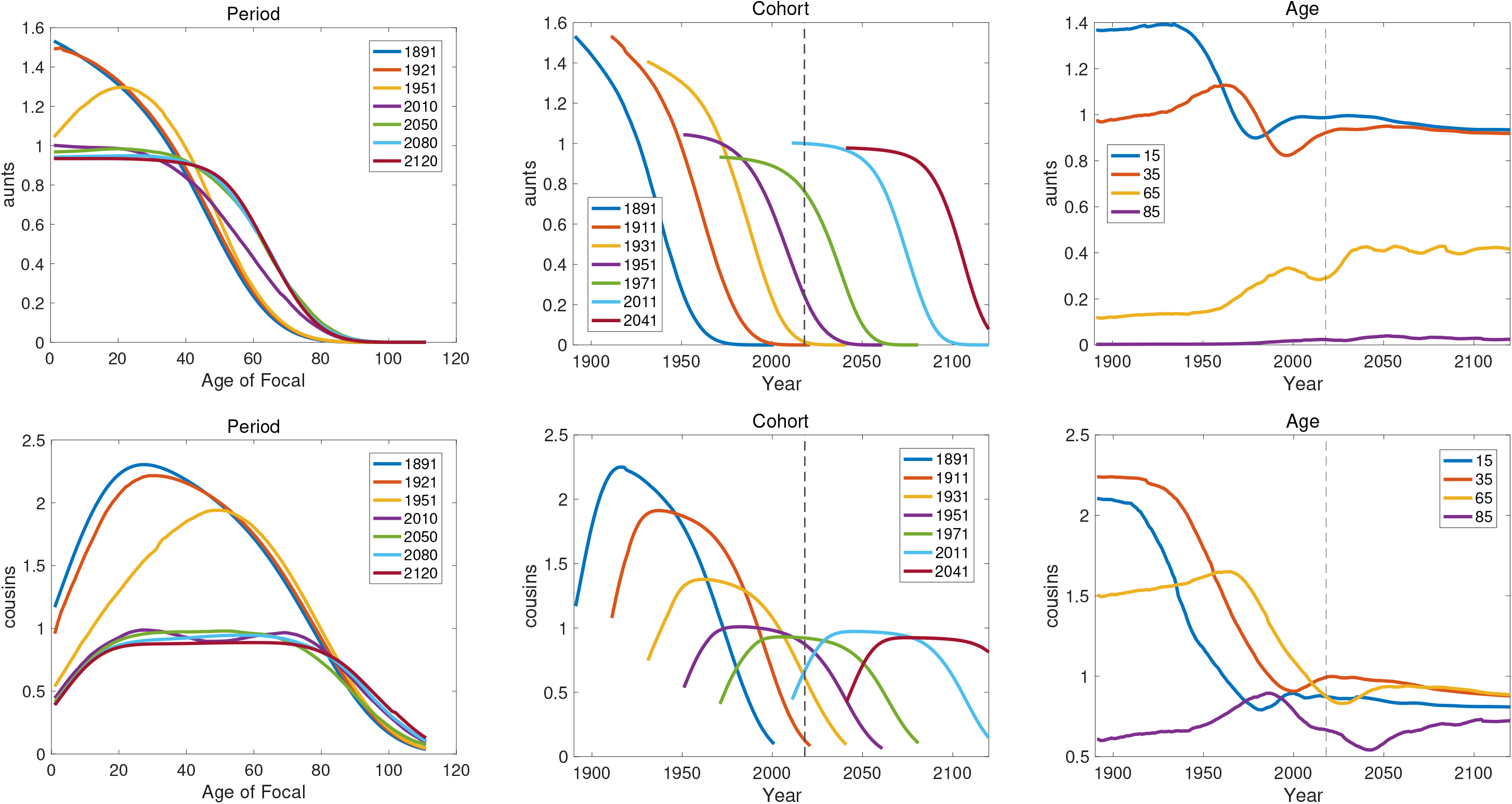
Period, cohort, and age results for the numbers of aunts and cousins of Focal; data from Sweden. Aunts older and younger than mother, and cousins through older and younger aunts as in Figure 1, have been combined.

For each kin type, the results are shown in a triptych of graphs in Figures 4-A to 4-E. The panels in these graphs show, from left to right,

1. Period results showing the numbers of kin, as the function of the age of Focal for a selection of years, from 1891 to 2120.
2. Cohort results, showing the numbers of kin of Focal over the lifetime of several cohorts, starting at a selection of years from 1891 to 2041.
3. Age results, showing the numbers of kin, in each year, of Focal at selected ages (15, 35, 65, and 85).

Several patterns are apparent in comparing these graphs. The period results fall into two general groups. In the early years (1891-1950), the numbers of kin are higher, and the peak gradually shifts to older ages of Focal. This is the reflection in kinship of the decline in fertility over that period (Figure 4). The later years (2010-2120) show lower abundances and much less of a decline over the lifetime of Focal, reflecting the very high survival and low fertility in recent and future Sweden.

The cohort results fall into several patterns. The abundance of one group of kin (daughters, granddaughters, great-granddaughters, and nieces^3^) is zero at Focal’s birth, increases, and then declines at later ages. Kin in a second category (mother, grandmothers, great-grandmothers, aunts) begins with positive numbers at Focal’s birth, and decline monotonically as Focal ages. The abundance of kin in a third category (sisters and cousins) begins with positive values at the birth of Focal, increases for a while, and then declines with the age of Focal.

## 4 Discussion

The analysis here retains the Focal-centric approach of Goodman, Keyfitz, and Pullum (1974) and follows Caswell (2019, 2020) in treating the kin of Focal as a population. Whereas in the time-invariant model dynamics unfold in relation to the age of Focal, in the model here, both Focal’s age and calendar time appear in the dynamics. This permits arbitrary temporal variation in the vital rates to be incorporated.

The analytical overhead is low; the main requirement is a sequence of survival and fertility matrices over some time interval, rather than a single matrix. The code for the time-varying case requires only minor changes from the time-invariant case (cf. Table 1 and Section A).

Because the form of the equations describing the dynamics is so similar to the time-invariant case, several aspects of the time-invariant theory apply directly.

### Properties of kin

The kinship structure vector **k**(*x, t*) contains the age structure of kin at age x of Focal at time t. This age structure provides the information needed to compute many interesting properties of the kin; e.g., dependency ratios, prevalence of health conditions (Caswell, 2019) and levels of unemployment (Song and Caswell, 2021)

### Extensions beyond age classification

As presented here, the model describes the age structure of kin as defined by age-specific demographic rates. The matrix **U**_*t*_ contains only probabilities of survival and aging. However, individual characteristics in addition to age may affect the demography, and the restriction to age classification is easily relaxed in this matrix framework. For example, Caswell (2019) extended the matrices and vectors to include the age structures of dead as well as living kin, and calculated Focal’s lifetime experience of the death of kin. Caswell (2020) extended the model to include, in addition to age, arbitrary stage structures with the possibility of movement among those stages, with an age×parity-dependent analysis as an example. In these multistate formulations, the model equations are unchanged; only the contents and block structures of the matrices change. Thus the time-varying model we present here applies directly to these more expansive individual state spaces, requiring only the substitution of the appropriate block-structured matrices and vectors.

### Time-varying rates in stochastic environments

The historical time-varying rates for Sweden are a fixed sequence. The future rates are generated by a deterministic projection process. But the world is actually stochastic, and there will be fluctuations in the rates due to that stochasticity. For example, the life expectancy in Sweden under 1989 rates dropped noticeably due to the influenza pandemic. The projected future life expectancy under 2020 rates shows no such decline due to the COVID-19 pandemic. Projections can be developed that incorporate environmental stochasticity (Lee and Tuljapurkar, 1994) and those or related methods could be used to create a stochastic time-varying kinship model. However, most of the theory of population growth in stochastic environments has focused on long-term ergodic properties of populations in stationary stochastic environments (e.g., Cohen, 1979; Tuljapurkar, 1990; Caswell, 2001). The kin of Focal develops over a time interval (Focal’s lifetime) that is short relative to the environmental changes. The properties of such a kinship network under environmental stochasicity is an open research problem.

### Comparisons with observed kinship data

Formal models, of kinship or anything else, are a way to deduce the implications of a set of demographic processes and parameters. These models provide a tool for demographers to examine how different components of population structure are connected to one another and to identify the effect of changing one component on population processes. Goodman, Keyfitz, and Pullum (1974) and Caswell (2019, 2020) have emphasized repeatedly that their models are not intended to predict the results of census data on the numbers of kin in a population. Such data will reflect processes operating in the real population but not included in the models; the models provide a background against which to compare the effect of those processes.^4^

Having incorporated time-varying demographic rates, the model we present here offers additional dimensions for comparisons with empirical measurements. Of the three types of results shown in Figures 4-A to 4-E, period results are perhaps the most easily measured empirically, from a single cross-setional measurement of the numbers of kin of individuals of various ages.

Measurements corresponding to cohort results would require longitudinal data on the numbers of kin of a set of individuals followed over their lives. Measurements corresponding to the age results would require information over time on the kin numbers of individuals at one or more selected ages. The challenges involved in obtaining both historical time series of demographic rates and empirical measurements of kin structure are obvious, but it is now possible to contemplate the calculations.

### Implications for multigenerational inequality

Social scientists have begun to extend the long-standing interest in parental influences on children to inequality over multiple generations and assess the influences of grandparents and other more distant kin in families’ long-term reproductive success and socioeconomic attainment (Mare, 2011; Song, 2021). Formal models of kinship complement existing empirical and simulation-based investigations of the demographic dynamics of households, lineages, and kin networks (LeBras and Wachter, 1978; Smith, 1987; Smith and Oeppen, 1993; Song, Campbell, and Lee, 2015). Social changes are often driven by various mechanisms of status transmission, not only from parents to offspring but also between other types of kinship relations, and by natural and cultural selections (Cavalli-Sforza and Feldman, 1981). Our formal models, in combination with multigenerational microdata of social mobility, allow researchers to expand the scope of future studies on social and demographic inequality among families over long timescales.

Two limitations of the model here (and other kinship models) are worth mentioning. First, the model uses female rates to calculate female kin (e.g., granddaughters) through female lineages (e.g., daughters of daughters). A full two-sex extension of the model removes this limitation and can incorporate differences between male and female rates (Caswell, 2021a). However, a crude approximation that accounts for both sexes under the hypothesis of equal male and female rates can be applied to the models presented here. The kin vector **k**(*x, t*) is multiplied by a constant that depends on the type of kin.^5^

The second limitation is the deterministic nature of the model. The age vectors **k**(*x, t*) are the mean age structures of kin. However, because the kin populations are small, demographic stochasticity will generate variance among individuals. That is, knowing the mean number of cousins of Focal at some age and time tells us nothing about the variance to be expected among Focal individuals experiencing the same demographic rates. Stochastic solutions of the dynamic equations provide information on the distribution of kin (Caswell, 2021b); more sophisticated branching process models may also be applicable (Pullum, 1982). We look forward to developments in these areas.

## 5 Acknowledgements

This research was supported by the European Research Council under the European Union’s Horizon 2020 research and innovation program, through ERC Advanced Grant 78819 to HC.

HC is grateful for helpful discussions with Silke van Daalen and the Theoretical Ecology Group at the University of Amsterdam and for the hospitality of the Max Planck Institute for Demographic Research.

XS is grateful to Thomas Coleman, Yongai Jin, Yue Qian, and Yu Xie for helpful discussions.

## A Time-invariant kinship calculations

For comparison with Table 1, this Appendix presents the components of the time-invariant model. Reproduced (with typos corrected) from Caswell (2019) under the terms of a CC-BY license.

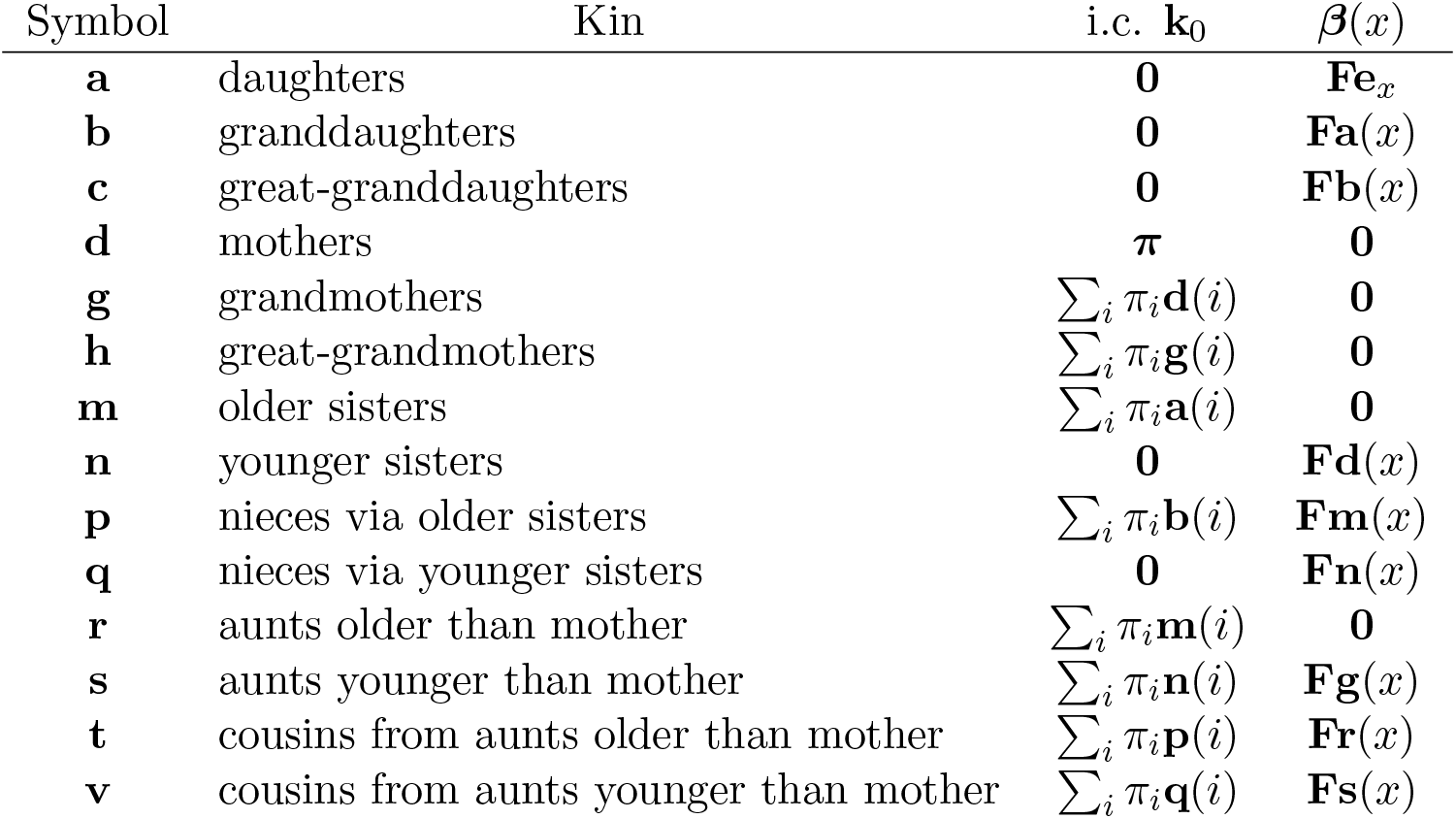

## Footnotes

1 For ease of comparison with the time-invariant case, the presentation here closely follows that of Caswell (2019), under the terms of a Creative Commons Attribution (CC-By) license.

2 The mortality rates were obtained from: https://www.statistikdatabasen.scb.se/pxweb/en/ssd/START__BE__BE0401__BE0401F/BefProgDodstal18/. The fertility rates were obtained from https://www.statistikdatabasen.scb.se/pxweb/en/ssd/START__BE__BE0401__BE0401F/BefProgFruktTot18/. As of this writing, the most recent description of the methods used in the projections is Statistics Sweden (2018), in Swedish only. An earlier version in English is available as Statistics Sweden (2012).

3 In principle, Focal could, at birth, have nieces through her older sisters, but the abundance in these data is extremely low.

4 See the essay by Nathan Keyfitz, “How do we know the facts of demography?” (Keyfitz and Caswell, 2005, Chap. 20) for a discussion of this.

5 Multiply daughters by 2, granddaughters by 4, great-granddaughters by 8, mothers by 2, grandmothers by 4, great-grandmothers by 8, sisters by 2, nieces by 4, aunts by 4, and cousins by 8. Goodman, Keyfitz, and Pullum (1974) first suggested this approximation.

